# Developmental exposure to perfluorooctane sulfonate (PFOS) and perfluorooctanoic acid (PFOA) selectively decreases brain dopamine levels in northern leopard frogs

**DOI:** 10.1101/623900

**Authors:** Rachel M. Foguth, R. Wesley Flynn, Chloe de Perre, Michael Iacchetta, Linda S. Lee, Maria S. Sepúlveda, Jason R. Cannon

## Abstract

Per- and polyfluoroalkyl substances (PFAS) are synthetic compounds that are a major public health concern due to widespread use, long environmental and biological half-lives, detection in most human plasma samples, and links to multiple adverse health outcomes. The literature suggests that some PFAS may be neurotoxic. However, there are major gaps in the literature with respect to how environmentally-relevant doses during development may influence the nervous system. To address this gap, we utilized a sentinel species, Northern leopard frogs (*Lithobates pipiens)* to determine the effects of developmental exposure to environmentally relevant perfluorooctane sulfonate (PFOS) and perfluorooctanoic acid (PFOA) on major neurotransmitter systems. Frog larvae at Gosner stage 25 were exposed to 10, 100, or 1000 ppb PFOS or PFOA for 30 days before neurochemical analysis. High performance liquid chromatography (HPLC) with electrochemical detection or fluorescent detection assays was used to measure neurotransmitter levels, which were normalized to protein levels in each sample. Dopamine (DA) decreased significantly in the brains of frogs treated with PFOA (1000 ppb) and PFOS (100 and 1000 ppb). Significant increases in DA turnover also resulted from PFOA and PFOS treatment. Neither PFOS, nor PFOA produced detectable alterations in serotonin (nor its metabolite), norepinephrine, gamma-amino butyric acid (GABA), glutamate, or acetylcholine. PFAS body burdens showed that PFOS accumulated relative to dose, while PFOA did not. These data suggest that DArgic neurotransmission is selectively affected in developmentally exposed amphibians and that PFAS should be evaluated for a potential role in diseases that target the DA system.

## 1. Introduction

Per- and polyfluoroalkyl substances (PFAS) are compounds that have been extensively used in manufacturing of surfactants, paper and packaging treatments, anti-static agents, corrosion inhibitors, insecticides, shampoos, and firefighting foams (Posner, 2012; Stahl *et al.*, 2011). PFAS are of significant concern due in part to the stable carbon-fluorine bond and, therefore, long half-life and potential for accumulation (Moody *et al.*, 2003). Perfluorooctane sulfonate (PFOS) and perfluorooctanoic acid (PFOA) have been extensively investigated for adverse health effects. While PFOS and PFOA have largely been phased out in favor of shorter chain length PFAS, they are ubiquitously detected in the global population (Jian *et al.*, 2018; Olsen *et al.*, 2009).

PFOS and PFOA both cross the blood-brain barrier, with PFOS generally accumulating 10-100x more in mammals than many other major PFAS (Maestri *et al.*, 2006). Studies in large mammals suggest that PFAS accumulate in the brain and are potentially neurotoxic. In polar bears, brain PFAS levels (including PFOS) were found to correlate with neurotransmitter alterations (Dassuncao *et al.*, 2019). Further, PFAS in North Atlantic pilot whales (*Globicephala melas*) were found to accumulate in brain, where only the liver had higher levels. Here, PFAS were strongly correlated with the high phospholipid content in the brain (Dassuncao et al., 2019). Laboratory studies in rodents on PFAS neurotoxicity have suggested neurobehavioral alterations. For example, mice exposed to various PFAS on post-natal day 10 showed that exposure increased spontaneous activity and increased proteins important for making synapses (Johansson *et al.*, 2009; Lee and Viberg, 2013). After pregnant dams were treated with PFOA, male offspring had significantly shorter latency to fall on the wire-hang test when they were 5-8 weeks old, suggesting a potential motor deficit (Onishchenko *et al.*, 2011). This implies that PFAS exposure may affect brain development. Exposures to invertebrates have shown that *C. elegans* have decreased motor function after 48 hour exposure to PFOS (Chen *et al.*, 2014). Finally, *in vitro* exposures suggest that PFAS may influence neuronal differentiation, where, in PC12 cells, PFOS promoted differentiation to cholinergic neurons at the expense of differentiation to dopaminergic (DA)rgic neurons, whereas other tested PFAS suppressed or enhanced overall differentiation (Slotkin *et al.*, 2008). This research suggests that developmental exposure could lead to a long-term deficit in the number of DA neurons.

In general, there is a lack of data on PFAS neurotoxicity and potential relevance to neurological diseases. Neurobehavioral alterations have been examined in a number of developmental and adult studies, with some behavioral endpoints suggesting the cholinergic system as a target (Johansson *et al.*, 2008). In vitro studies have suggested glutamatergic signaling as a target (Liao *et al.*, 2009). However, environmental relevance of the tested doses is generally lacking and to our knowledge no studies measured neurotransmitter levels from multiple systems in response to PFAS. The primary goal of this report was to determine the effects of developmental PFAS exposure at doses which include those detected in environmental samples in a sentinel species. The long-term goal is to prompt specific research into how exposure to PFAS during development may affect specific neurotransmitter systems and influence neurological disease.

## 2. Methods

### 2.1 Animals, nominal exposure, and measured water and sediment PFAS levels

All animal studies were approved by Purdue Animal Care and Use Committee standards. The Northern leopard frog (*Lithobates pipiens*) was chosen as a relevant sentinel species (Hoover *et al.*, 2017). Eggs were collected and reared to Gosner stage 25 in outdoor tanks prior to the start of the experiment. Treatment tanks were prepared with 75 L of aged well water (containing algae and zooplankton) and 10 kg of sediment spiked with PFOS or PFOA to achieve nominal sediment concentrations of 0, 10, 100, or 1000 ppb. We chose to spike sediments, instead of simple addition of chemicals to the water to represent a more ecologically relevant exposure scenario. Tadpoles unintentionally consume sediment while grazing for algae and detritus. Thus, spiking sediments provided more realism in incorporating all pertinent routes of exposure amphibian larvae would likely experience in the environment. The doses were selected to include concentrations detected at contaminated sites (Moody *et al.*, 2003). Measured water and sediment concentrations, along with body burdens were determined by tandem mass spectrometry from representative samples as previously reported (Hoover *et al.*, 2017). Exposure lasted 30 days and tadpoles were sacrificed immediately using tricaine (MS-222). For this short initial report on neurotoxicity, 6-14 frogs/treatment (PFOA or PFOS at 0 to1000 ppb) were randomly selected from 32 treatment tanks (~20 tadpoles housed/tank) for detailed neurotransmitter analysis (81 total samples analyzed for each neurotransmitter), where the majority of the animals were devoted to a long-term, non-neuronal study on systemic ecotoxicity. Brains were removed, frozen in liquid nitrogen, and stored at −80°C until processed.

### 2.2 Neurotransmitter Quantification

Six neurotransmitter systems were assessed. The majority of neurotransmitters were analyzed by high performance liquid chromatography (HPLC) with electrochemical detection as we have previously reported (Agim & Cannon, 2018; Cannon et al., 2009; Horowitz et al., 2011; Wirbisky, Weber, Lee, Cannon, & Freeman, 2014; Wirbisky et al., 2015). Neurotransmitter levels in the samples were quantified by measuring the area under the curve and comparing it to a standard curve. Neurotransmitter levels were then normalized to protein levels in the sample (ng neurotransmitter/mg protein). DA turnover was calculated as [(DOPAC + HVA)/DA where DOPAC and HVA refer to DA metabolites 4-dihydroxyphenylacetic acid and homovanillic acid, respectively] (Agim and Cannon, 2018).

Acetylcholine (ACh) was quantified using the Invitrogen Molecular Probes Amplex ACh/Acetylcholinesterase Assay Kit (A12217). Briefly, samples were diluted 1:1 in the reaction buffer. Samples and standards were incubated in 200 μM amplex red, 1 U/mL horseradish peroxidase, 0.1 U/mL choline oxidase and 0.5 U/mL acetylcholinesterase 1:1 reaction buffer. Concentrations were determined with fluorescence spectroscopy (530 nm excitation and 590 nm emission). Acetylcholine levels were normalized to protein levels in the sample (ng neurotransmitter/mg protein).

### 2.4 Statistical Analysis

Differences in bioaccumulation of PFAS between treatments were tested using linear regression on log10-transformed PFAS body burdens and comparisons of estimated marginal means post-hoc analysis. Neurotransmitters were analyzed by one-way ANOVA with Sidak’s post-hoc test to compare all treatment groups within an exposure (PFOA or PFOS) and to controls. Whole brain analyses of lower order species (i.e., zebrafish) produce considerably more variable neurochemistry data than micro-dissected brain analysis in rodents (Dukes *et al.*, 2016; Lee *et al.*, 2015). Thus, outlier analysis was conducted. Statistical analyses were conducted both before and after outlier analysis. Importantly, outlier analyses for raw neurotransmitter data had no effect on statistical significance, where no new significant comparisons were revealed, nor were any significant comparisons eliminated after outlier analysis. Outliers were identified using stringent ROUT analysis (Q = 0.1%) (Motulsky and Brown, 2006). Outlier analyses resulted in no more than 3 samples/group being excluded, with the following total exclusions/endpoint: DA (0/81 data points omitted), DOPAC (7/81 data points omitted), HVA (0/81 data points omitted), 5-HT (0/81 data points omitted), 5-HIAA (0/81 data points omitted), NE (1/81 data points omitted), DA turnover (9/81), 5-HT turnover (1/81 data points omitted), glutamate (0/81 data points omitted), GABA (11/81 data points omitted), and acetylcholine (0/81 data points omitted). All data were expressed as mean ± standard error of the mean (SEM) and *p* < 0.05 used to identify significant differences.

## 3. Results

### 3.1 Water, sediment, and body burden levels

Quantified PFAS exposure and body burden levels are shown in Table 1 and Figure 1.

**Table 1.** PFAS applied and measured doses and total body burden. Measured concentrations include sediment and water analysis, along with whole body burden determination at the end of the 30-day exposure (*n* = 4-5). Control values = (PFOA + PFOS).

| Exposure | Nominal sediment concentration (ppb) | Measured water concentration at initiation (ppb ± SEM) | Measured water concentration at completion (ppb ± SEM) | Measured sediment concentration at initiation (ppb ± SEM) | Measured sediment concentration at completion (ppb ± SEM) | Body burden (ppb ± SEM) |
| --- | --- | --- | --- | --- | --- | --- |
| Control | 0 | 0.000 ± 0.000 | 0.000 ± 0.000 | 0.102 ± 0.060 | 0.0778 ± 0.0768 | 27.7 ± 9.2 |
| PFOA | 10 | 3.22 ± 0.12 | 2.58 ± 0.17 | 18.9 ± 1.0 | 7.61 ± 0.40 | 205 ± 26 |
| PFOA | 100 | 8.34 ± 0.73 | 6.33 ± 0.13 | 42.0 ± 1.6 | 14.3 ± 1.1 | 161 ± 17 |
| PFOA | 1000 | 76.911 ± 2.603 | 55.379 ± 0.575 | 423 ± 22 | 140 ± 15 | 169 ± 24 |
| PFOS | 10 | 0.0812 ± 0.0390 | 0.0326 ± 0.0295 | 7.89 ± 0.34 | 5.94 ± 0.67 | 196 ± 41 |
| PFOS | 100 | 0.771 ± 0.078 | 1.15 ± 0.13 | 89.1 ± 4.7 | 45.6 ± 2.8 | 853 ± 46 |
| PFOS | 1000 | 16.4 ± 0.4 | 14.8 ± 0.3 | 739 ± 29 | 456 ± 45 | 4500 ± 660 |

**Figure 1.**
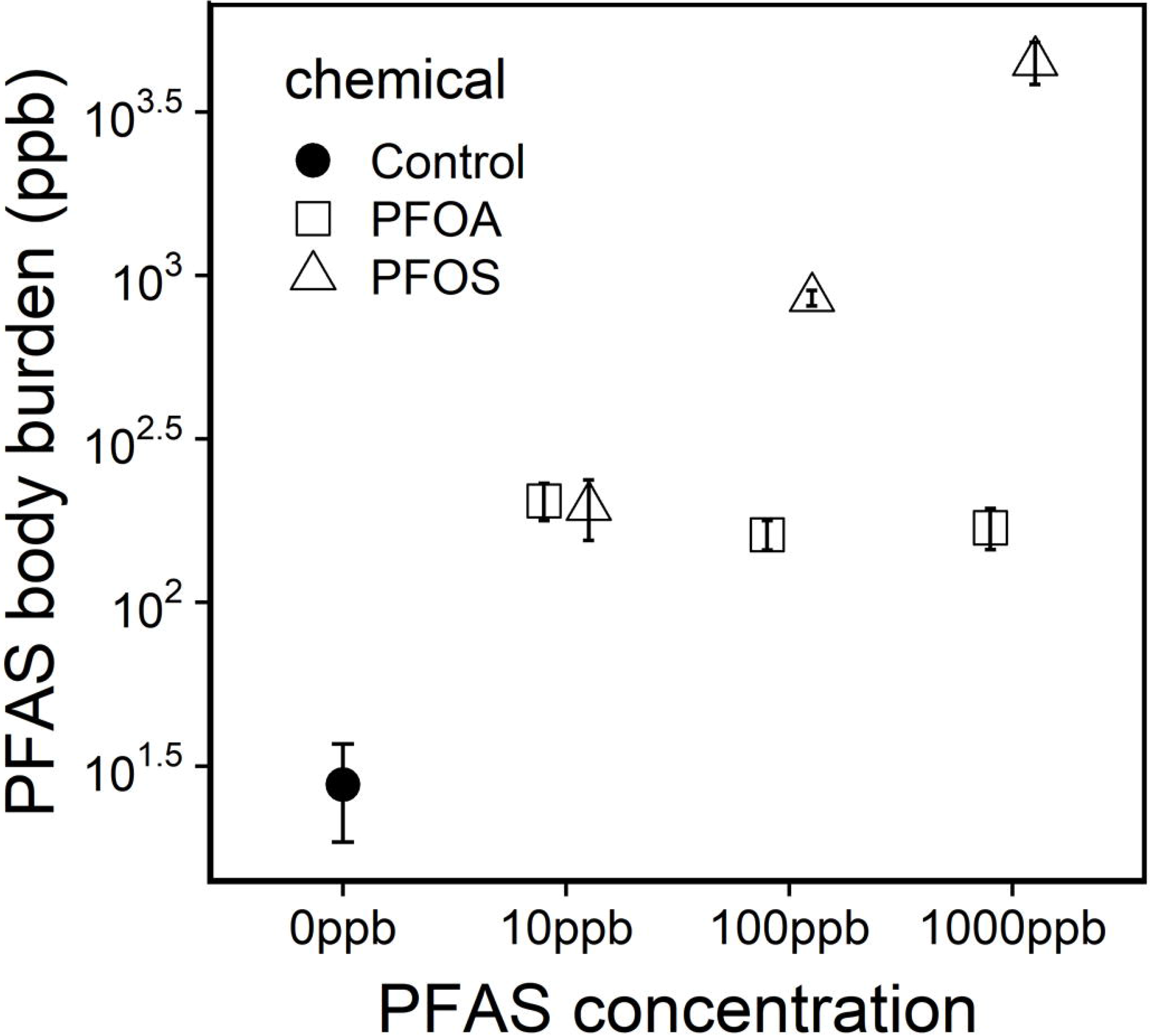
Total body burden in PFAS-treated frogs 30 days after treatment (mean +/− SEM). A regression shows that all PFAS treatments have greater body burdens than controls. Post-hoc analysis showed that: 1) the two highest PFOS treatment were significantly greater than the low PFOS treatment and all PFOA treatments, 2) the 1000ppb PFOS treatment was greater than the 100ppb PFOS, and 3) that none of the PFOA treatments differed from each other. Note, y-axis is log_10_-transformed.

### 3.2. Neurotransmitter levels

Quantification of catecholamine neurotransmitters showed a significant decrease in DA in frogs treated with 1000 ppb PFOA and 100 or 1000 ppb PFOS, (3.61 ± 0.61 ng/mg protein, 1.62 ± 0.18 ng/mg protein, 2.41 ± 0.15 ng/mg protein, 2.09 ± 0.24 ng/mg protein for the control, 1000 ppb PFOA, 100 ppb PFOS, and 1000 ppb PFOS, respectively) (Figure 2A), but no significant changes in the DA metabolites DOPAC or HVA (Figure 2B, C). DA decreases, relative to metabolite levels led to a significant increase in DA turnover [(DOPAC + HVA)/DA] for both 1000 ppb PFOS and PFOA (0.332 ± 0.051, 1.22 ± 0.20, 0.693 ± 0.085, 1.02 ± 0.10 for the control, 1000 ppb PFOA, 100 ppb PFOS, and 1000 ppb PFOS, respectively) (Figure 2D). Neither PFOS nor PFOA caused significant changes in norepinephrine (NE) levels (Figure 2E). Neither serotonin (5-HT) nor its metabolite 5-hydroxyindoleacetic acid (5-HIAA) were significantly affected by treatment with either PFAS, leading to no changes in 5-HT metabolism (Figure 2F - H).

**Figure 2.**
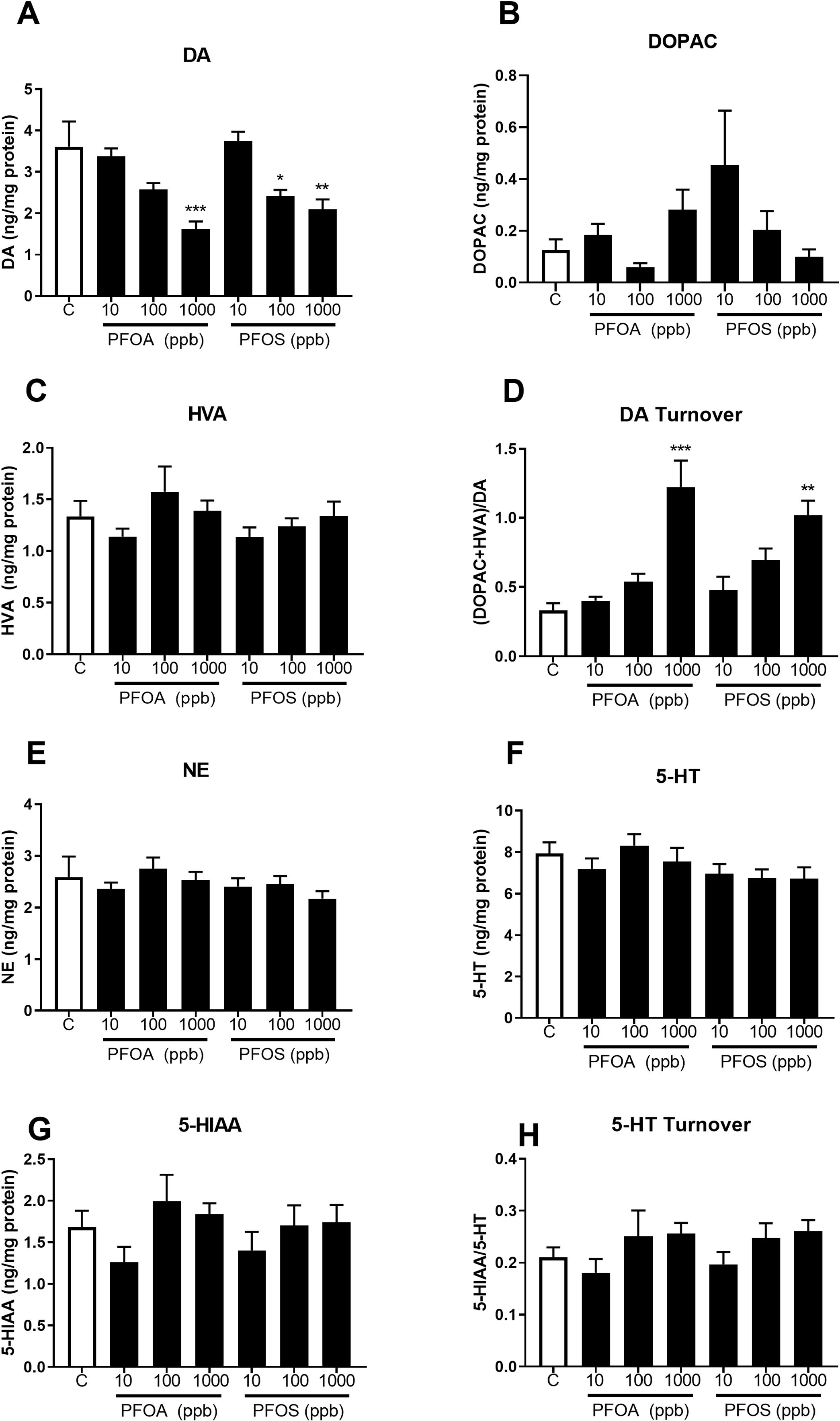
Developmental exposure to PFOA or PFOS selectively decreases dopamine levels. Northern leopard frog (*Lithobates pipiens*) tadpoles at Gosner stage 25 were exposed to PFOS or PFOA for 30 days. Whole brain monoamines and major metabolites were quantified using HPLC with electrochemical detection and normalized to amount of protein per sample: Dopamine (DA) (***A***); 3,4-dihydroxyphenylacetic acid (DOPAC) (***B***); homovanillic acid (HVA) (***C***); DA turnover [(DOPAC + HVA)/DA] (***D***); norepinephrine (NE) (***E***); serotonin (5-HT) (***F***); 5-hydroxyindoleacetic acid (5-HIAA) (***G***); 5-HT turnover (5-HIAA/5-HT) (***H***). Data was analyzed using one-way ANOVA followed by Sidak’s post hoc test. *p<0.05, **p<0.005, ***p<0.0001; n=6-14/group.

The amino acid neurotransmitters glutamate and gamma-aminobutyric acid (GABA) were measured and no significant differences were found in treatment groups versus the control for either glutamate or GABA (Figure 3A, B). In addition, there was no significant change in ACh levels in any treatment group compared to the control (Figure 3C).

**Figure 3.**
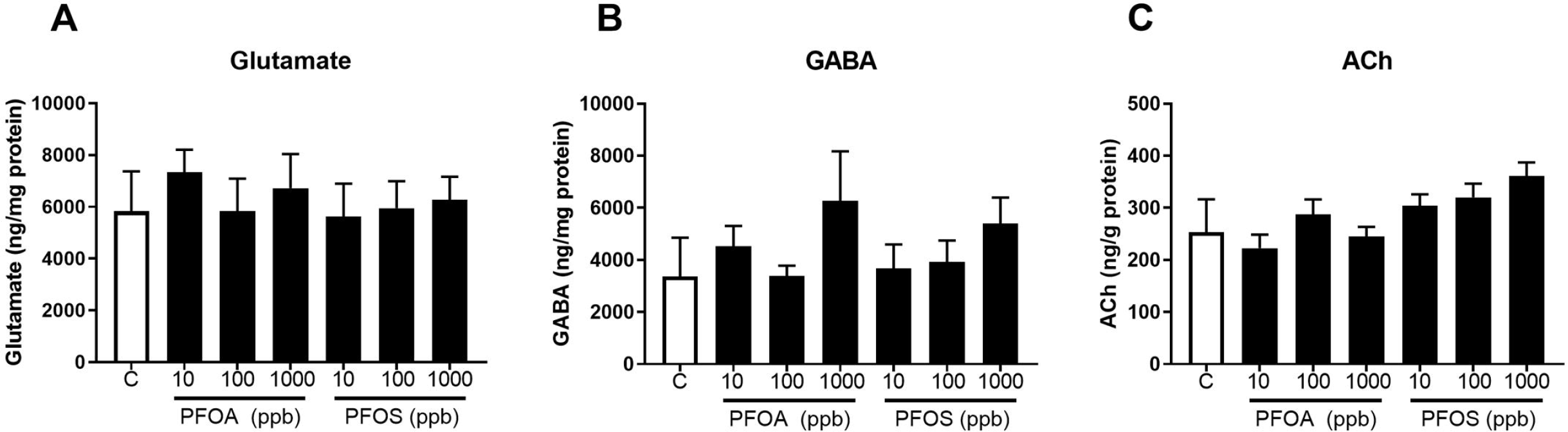
Developmental exposure to PFOA or PFOS does not affect amino acid or cholinergic neurotransmitter levels. Northern leopard frog (*Lithobates pipiens*) tadpoles at Gosner stage 25 were exposed to PFOS or PFOA for 30 days. Whole brain amino acid neurotransmitters were quantified using HPLC with electrochemical detection: glutamate (***A***); and gamma-aminobutyric acid (GABA) (***B***). Whole brain acetylcholine was quantified using the Invitrogen Molecular Probes Amplex Acetylcholine/Acetylcholinesterase Assay Kit with fluorometric detection. Neurotransmitters were normalized to amount of protein per sample. Data was analyzed using one-way ANOVA followed by Sidak’s post hoc test; n=6-14/group.

## 4. Discussion

There are major gaps in the literature related to PFAS neurotoxicity. Here, we have shown that developmental exposure within the concentration range documented at contaminated sites produce selective total brain DA depletion in an amphibian sentinel species. Further, in general, we observed that PFAS exposure increased DA turnover, which is typically a compensatory mechanism in response to DArgic neurotoxicity and DA depletion; and an additional metric of damage to DArgic function (Sossi *et al.*, 2002). This short report is expected to prompt research on the neurochemical and neuropathological effects of PFAS on the brain. Further, our findings suggest that the potential role of PFAS in neurological diseases affecting DArgic neurotransmission should be examined.

DA depletion in the frog brain could have several implications, especially given that there are some significant advantages of specific frog species as a model. Leopard frogs are unique in that they have neuromelanin-containing DArgic neurons in the ventral motor neurons which could be related to the DArgic neurons in the nigrostriatal DA system that are affected in Parkinson’s disease (PD). Importantly, these pigmented neurons have been shown to bind PD-relevant toxicants, where high-neuromelanin expressing species exhibit heightened sensitivity with respect to low-neuromelanin expressing species (Karlsson *et al.*, 2009; Lindquist *et al.*, 1988; Sokolowski *et al.*, 1990; Spencer *et al.*, 2004). Taken together, the literature suggests that, with respect to neuromelanin expression and toxicant binding to neuromelanin, certain amphibian species are more representative of human DArgic neurons than low-expressing rodent models. Thus, the potential relevance of DA depletion in PFAS-treated amphibians to PD should be further examined. An additional common DArgic pathway in mammals is the reward system with DArgic neurons extending from the ventral tegmental area to the ventral striatum (Lammel *et al.*, 2014). Frogs have a similar DArgic pathway for reward that contains DArgic neurons in the anteroventral tegmental area, thus the DA depletion that we observed could potentially result in suppression of the reward pathway (O’Connell *et al.*, 2010). DArgic neurons are also important for the hypothalamic function and multiple neuroendocrine axes that control well-conserved behaviors such as metabolic rate stress, digestion, and mood (Hu *et al.*, 2008). Thus, neuroendocrine function in response to PFAS should also be tested.

Another important aspect of these findings potentially relates to developmental origins of adult health and disease (DOHaD). It will be important to examine whether developmental PFAS exposures result in increased susceptibility to neurodegenerative diseases. PD, for example is a disease of aging, where, once the nigrostriatal DA reserve is depleted (i.e., loss of DA neurons reaches a symptomatic threshold), the onset of clinical symptoms appear (Tabbal *et al.*, 2012). Testing whether a developmental PFAS exposure may lower such reserve and increase susceptibility later in life is of importance.

Relationships between PFAS treatments and body burden show log-linear bioaccumulation of PFOS, whereas PFOA did not, which is consistent with the bioaccumulative properties of PFOS, especially in the brain (Maestri *et al.*, 2006). Differences between nominal and measured sediment PFAS levels are due to redistribution of the PFAS between sediment and water, and to a lesser part from a mass balance perspective, to uptake by algal communities and the frogs. Mass balance checks (water, sediment, and frogs) are generally within 15% of the nominal mass added to the system and calculated sediment-water partition coefficients are in the range reported in the literature (Higgins and Luthy, 2006). PFAS levels observed in control animals were low and to be expected, given the ubiquitous global distribution of PFAS. We hypothesize that PFAS detected in control animals could be derived from the natural water source used to fill tanks, algae, zooplankton and/or leaching from the plastic.

There are some notable limitations in the present study. Neurochemical quantification is a frequent endpoint studied for effects of compounds on the brain in rodents, but has also been measured in aquatic species, both in whole zebrafish larva and brains (Cannon *et al.*, 2009; Dukes *et al.*, 2016; Milanese *et al.*, 2012; Wirbisky *et al.*, 2014; Wirbisky *et al.*, 2015). However, quantification in aquatic species is more difficult and more variable than specific region neurotransmitter quantification in rodents. Clearly, our samples exhibited increased variability relative to micro-dissected rodent studies. Indeed, GABA and ACh analyses exhibited potential nonsignificant trends resulting from PFAS treatment, suggesting further examination. Future studies will require an increased number of animals/group, along with sub-regional analysis.

In summary, our findings suggest developmental PFAS exposure in a sentinel species at environmentally-relevant doses produce selective DA depletion. These findings imply that the role of PFAS in neurological diseases involving DArgic neurotransmission should be evaluated.

